# Estimating Species-specific U.S. Waterfowl Harvest

**DOI:** 10.1101/2024.07.27.603620

**Authors:** Ben C. Augustine, J. Andrew Royle

## Abstract

The U.S. Fish and Wildlife Service monitors species-specific waterfowl (ducks, seaducks, geese, and brant) harvest through two hunter surveys, one that estimates the total harvest for each waterfowl group, and a second that estimates the species composition of each waterfowl group. Point estimates for species-specific harvest can be computed by multiplying the estimated total harvest by the estimated proportion of the total harvest of each species. However, to date, no uncertainty estimates have been available. Here, we combine these two data sources to provide species-specific harvest estimates at the state and flyway level while characterizing the uncertainty via Bayesian estimation. We take a similar approach to Smith *et al*. (2022), providing both estimates that treat yearly data as independent and estimates that share information across years via a random walk process. We then discuss the advantages and disadvantages of each approach.

## Introduction

The U.S. Fish and Wildlife Service (USFWS) requires accurate and precise estimates of state and flyway-level species-specific harvest of waterfowl to assess species status and trends, and to inform species management by the establishment of annual harvest regulations. These estimates are obtained through two sets of hunter surveys. The Waterfowl Harvest Survey (WHS) is used to estimate the total harvest at the state level for four groups of waterfowl–ducks, seaducks, geese, and brant. Then, the Parts Collection Survey (PCS) is used to estimate the species composition of the total harvest for each of these four species groups, with the exception of brant, for which only one species occurs in each state. The WHS asks a sample of hunters in each state and year how many birds they harvested (separate hunter samples for each species group), and the PCS uses a smaller sample of hunters in each state that send in bird parts such as wings and tails for agency biologists to identify to species and age class (when possible). Together, these data can be used to estimate species-specific total harvest, with the uncertainty of each data source accounted for.

Recently, Smith *et al*. (2022) used analogous data from the Canadian Wildlife Service to estimate species-specific total harvest using hierarchical models and Bayesian estimation. Smith *et al*. (2022) used temporal random walk models for both the total harvest and species composition data to improve the accuracy and precision of species-specific total harvest estimates. Here, we take a similar approach using data from the USFWS to 1) describe and apply hierarchical models to estimate species-specific total harvest, 2) assess the precision of species-specific total harvest at the state and flyway level achieved with the current WHS and PCS protocols (e.g., yearly sample sizes), and 3) compare the estimates between models that do and do not consider temporal correlation across years.

## Methods

### Methods - Data Structure

We develop models independently for each of four species groups: ducks, sea ducks, geese, and brant. For each data set, there are two sources of data–data related to the total harvest (WHS) and data related to the species composition of the harvest (PCS), with the exception of 1) brant, which has no species composition data because no state or flyway has more than one species, and 2) geese, for which we will also use data on age class (subadult vs. adult; data also from the PCS survey) to produce species by age class total harvest estimates. The total harvest data are structured by region and year, and the PCS data are structured by region, year, and species. We describe the model in terms of general ”regions”, but in this specific case, the regions are U.S. states. For this analysis, we consider 17 years of data for all species groups (2003 - 2019).

For each species group, the total harvest data consist of the region by year point estimates and standard errors of the total harvest computed from the hunter-level data for all *R* regions and *Y* years. These estimates are justified based on the survey methodology that follows (Dillman *et al*., 1978). We define 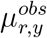 and 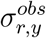 to be the point estimate and standard deviation, respectively, for the total harvest in region *r* and year *y*. Then, the PCS data consist of the region by year by species counts of submitted birds from the PCS. We define *parts*_*r,y,s*_ to be the number of submitted species parts in region *r* and year *y* for species *s*, and 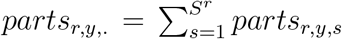 to be the total number of submitted parts in region *r* and year *y* summed across the *S*^*r*^ species in region *r*. Finally, the age data for geese in region *r*, year *y*, and age class *a* is structured as *age*_*r,y,s,a*_ for *a* = 1 …, 2, where *a* = 1 indicates ”adult” and *a* = 2 indicates ”subadult”. The total number of observations summed across age classes is 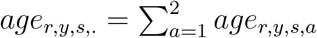.

### Methods - Models

To estimate region by year species-specific harvest for each species group from these two data sources, we model the region by year total harvest and species composition data separately, and then combine the estimates to derive species-specific harvest estimates (described below). These analyses are done independently across species groups. For geese, we additionally model the region age class composition and combine these with the species-specific harvest estimates to produce species and age class-specific harvest estimates. For both the total harvest model and PCS models (species composition and age class), we consider two general model structures, one that treats all regions and years as independent (independence model), and a second that shares information across time within each region using a temporal random walk (time model). Finally, these region (state) posterior distributions can be aggregated to larger spatial regions (e.g., flyways) by adding the posterior distributions (Smith *et al*., 2022) for regions contained in the larger spatial regions (e.g., adding posterior distributions of states in a particular flyway).

### Methods - Total Harvest Models

For the total harvest models, we assume that for region *r* and year *y*, 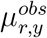, is normally distributed:

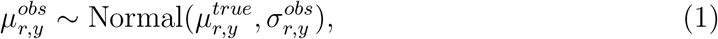

where 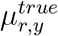 is the true harvest which we want to estimate. This model is used for both the independence and time models, with the differences between these two models being the structure of their prior distributions. For the independence model, we use a relatively diffuse prior of 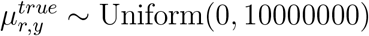 for each region and year that is meant to reflect the absence of prior information. This is uninformative in the sense that it is uniform over plausible values of the total harvest.

For the total harvest time model, we share information across years within a region using a random walk process (Lawler & Limic, 2010) to model the temporal correlation through time within a region on the log scale. We use the log scale so that the temporal correlation is with respect to the proportional, rather than absolute, change between years. First, we provide a prior distribution for the expected harvest in the first year of 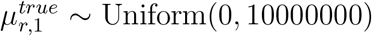, matching that in the independence model. Then, we assume the total harvest within a region follows a random walk on the log true harvest:

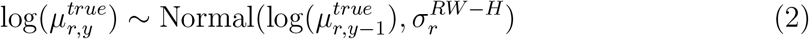

where 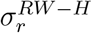 is the random walk standard deviation for region *r* (“H” indicating “harvest” model).

### Methods - PCS Model for Species Composition

For the PCS species composition data, we use Poisson regression (Dobson & Barnett, 2018) with the species proportions computed as derived parameters. This differs from the Dirichlet-multinomial approach of (Smith *et al*., 2022) that directly models the species proportions. We opted for Poisson regression because we were unable to get the Dirichlet-multinomial model to converge reliably with a random walk through time.

We assume

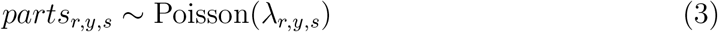

where *λ*_*r,y,s*_ is the expected number of submitted birds in region *r* in year *y* for species *s*. To account for different numbers of hunters submitting birds across regions and years, we use an offset (Dobson & Barnett, 2018), assuming

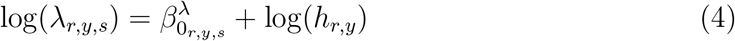

where *h*_*r,y*_ is the number of hunters in region *r* and year *y*. Therefore, we are modeling the rate of submitted birds per hunter, and 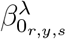 is the log rate of submission.

For the time model, we assume the log rate of submission follows a random walk through time:

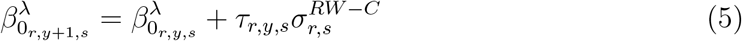

where

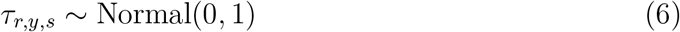

and 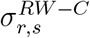 is the random walk standard deviation for region *r* and species *s* (”C” indicating ”PCS species composition” model). Note, this is a non-centered random effects parameterization (Papaspiliopoulos *et al*., 2003) that was necessary to achieve adequate Markov chain Monte Carlo (MCMC) mixing for many region-species. For both models, the species proportions are derived following

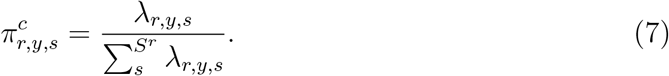

For the independence model, we specify the weakly-informative prior distributions, 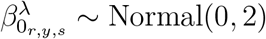 for all *r, y*, and *s*. For the time model, we used a weakly informative prior on the log submission rate for the first year of every species-region, 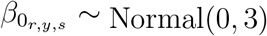 and for the random walk standard deviations, 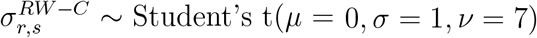.

### Methods - PCS Models for Age Class Composition

We use the same model and methods for the PCS age class data for geese that we do for species composition except the proportion of interest is with respect to age class for each species. To accommodate missing age class data for birds where age could not be determined, we assumed age class assignments were missing completely at random (Little & Rubin, 2019) and discarded the missing data. Similar to the PCS species composition model, we assume

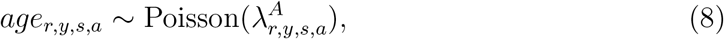

and model the log rate of submission by age

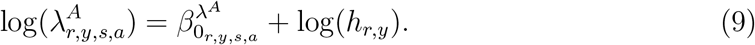

With the age class proportions derived following

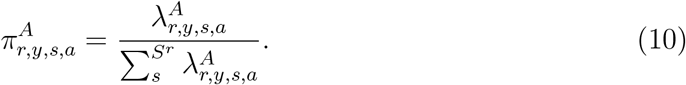

### Methods - Deriving Species-specific Harvest

The posterior distributions for region and species-specific harvest, 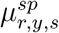 are obtained by multiplying the posterior samples of total harvest by the posterior samples of PCS species proportions for every MCMC iteration *i*:

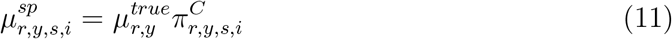

The posterior distributions for region, species, and age-specific harvest for goose, 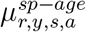 are obtained by multiplying the species-specific harvest samples by the PCS age class proportion samples for every MCMC iteration *i*:

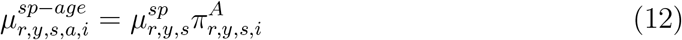

Finally, the flyway-level species and species by age harvest posterior distributions are obtained by adding the posterior distributions for the states in a particular region. For flyway-level duck posterior distributions, we combine the state level duck and sea duck posterior distributions because sea ducks are included in the duck category in some states.

### Methods - Parameter Estimation

We fit all models using the Nimble MCMC software (de Valpine *et al*., 2022, Version 0.13.1) in Program R (R Core Team, 2021, Version 4.0.5). We used default MCMC samplers for all parameters. We used posterior means as point estimates and 95% highest posterior density (HPD) intervals for interval estimates. For measures of estimate precision, we used the posterior coefficient of variation (CV), defined as the posterior standard deviation divided by the posterior mean multiplied by 100. We assessed convergence using the Gelman-Rubin statistic, ensuring that the 95% CI upper bound was less than 1.1.

## Results

### Results - State-Level Species-Specific Harvest Estimates

The independence and time models generally produced similar point estimates (Figure 1) for each of the four groups of species, except for species that were rare in the PCS data, which were generally estimated lower in the time models. These cases occurred mainly for brant, though there are examples for all species groups. Then, for all species groups, the posterior standard deviations were lower in the time model (Figure 2), with species-years with the largest standard deviations seeing the largest absolute reduction in the time model. Finally, the CVs for the time model were lower for ducks, seaducks, and geese, on average; however, the CVs were substantially higher for most brant and some sea ducks and geese (Figure 3; see Discussion).

**Figure 1:**
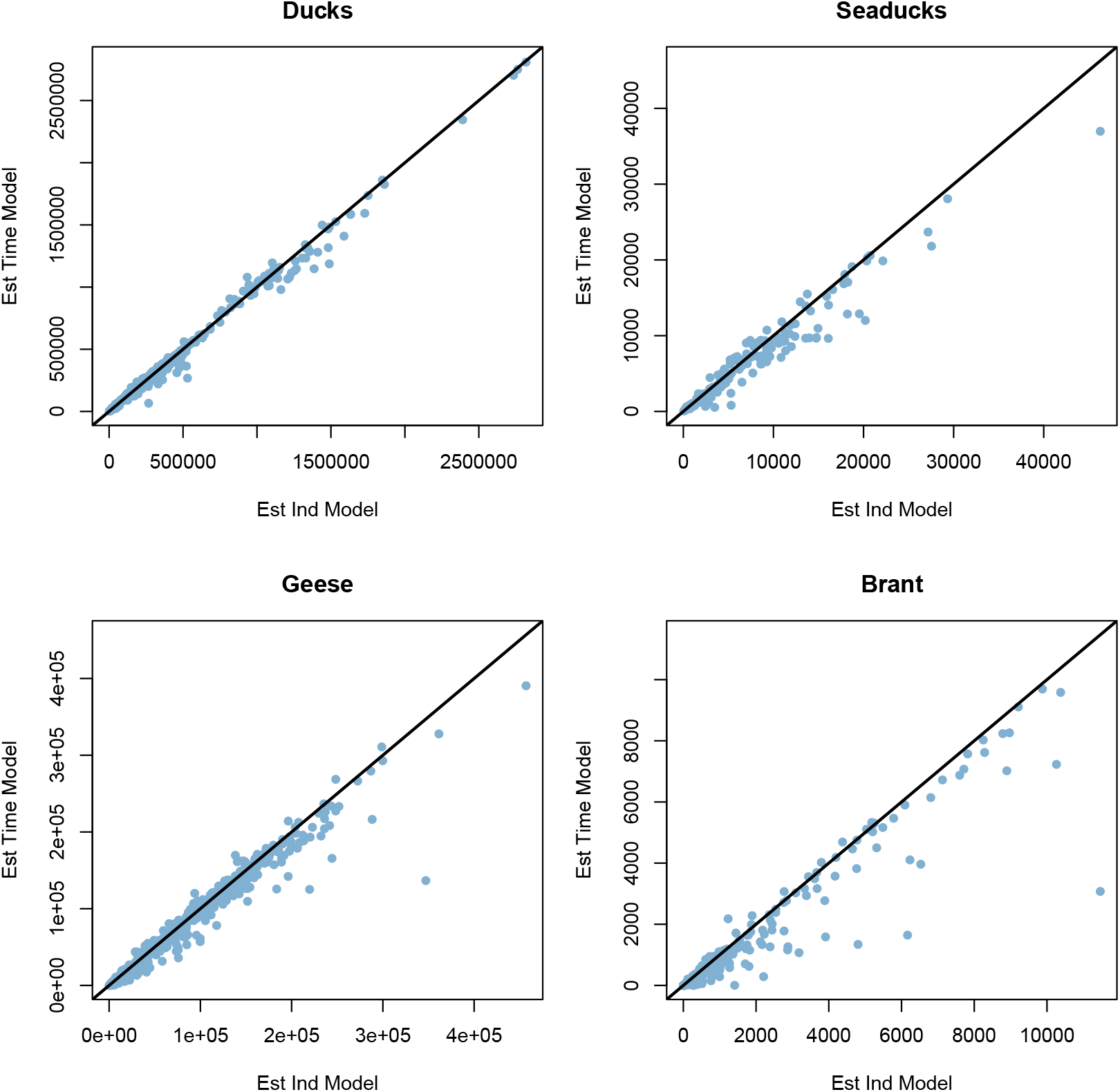
Comparison of state by species by year point estimates for each species group (ducks, seaducks, geese, and brant). Estimates from the independence and time models are on the x and y axes, respectively. The black line indicates equality between the two models.

**Figure 2:**
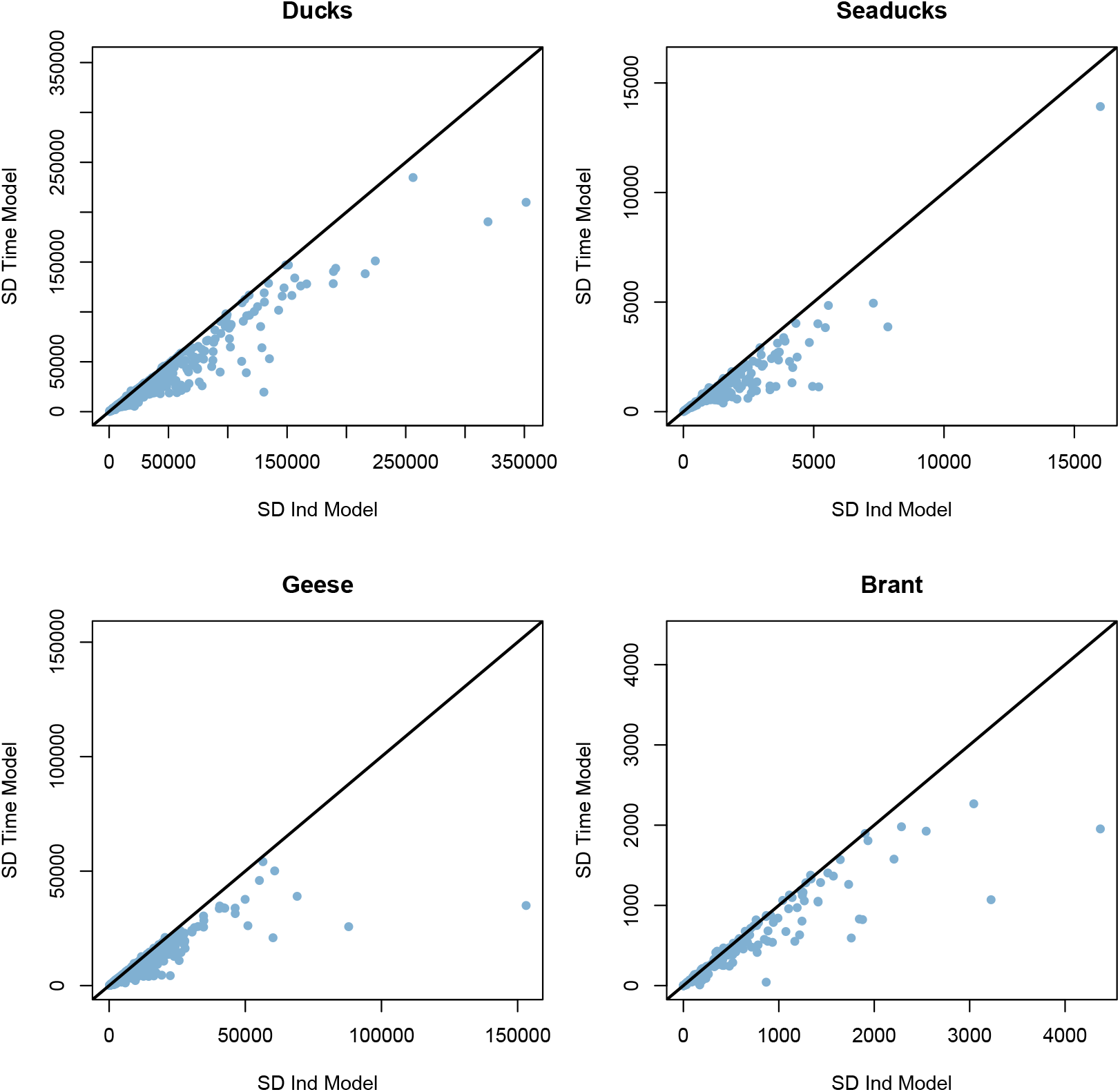
Comparison of state by species by year posterior standard deviations for each species group (ducks, seaducks, geese, and brant). Estimates from the independence and time models are on the x and y axes, respectively. The black line indicates equality between the two models.

**Figure 3:**
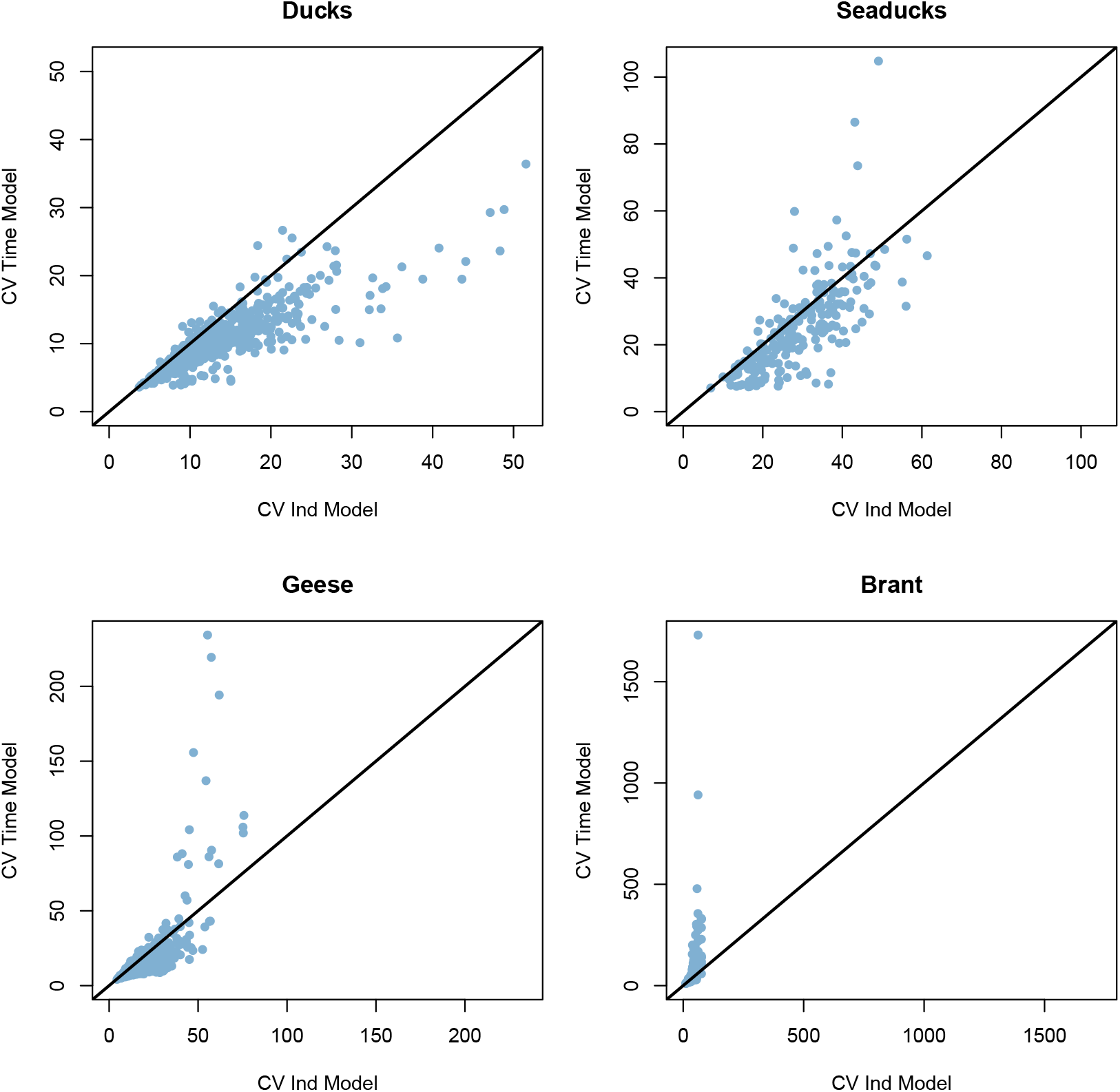
Comparison of state by species by year coefficients of variation for each species class. Estimates from the independence and time models are on the x and y axes, respectively. The black line indicates equality between the two models.

The degree to which the time model estimates differed from the independence model depended on the degree of temporal correlation in the time series (how similar are harvest levels through time) and number of PCS observations for a particular state-species. For example, Figure 4 shows an infrequently harvested species, Figure 5 shows a commonly harvested species, and the temporal model estimates differ more from the independence model for the infrequently harvested species. Figure 6 shows a more rarely harvested species where the time model that shares information across time estimates a much lower total harvest than the independence model that does not share information across time and is limited by yearly sample sizes.

**Figure 4:**
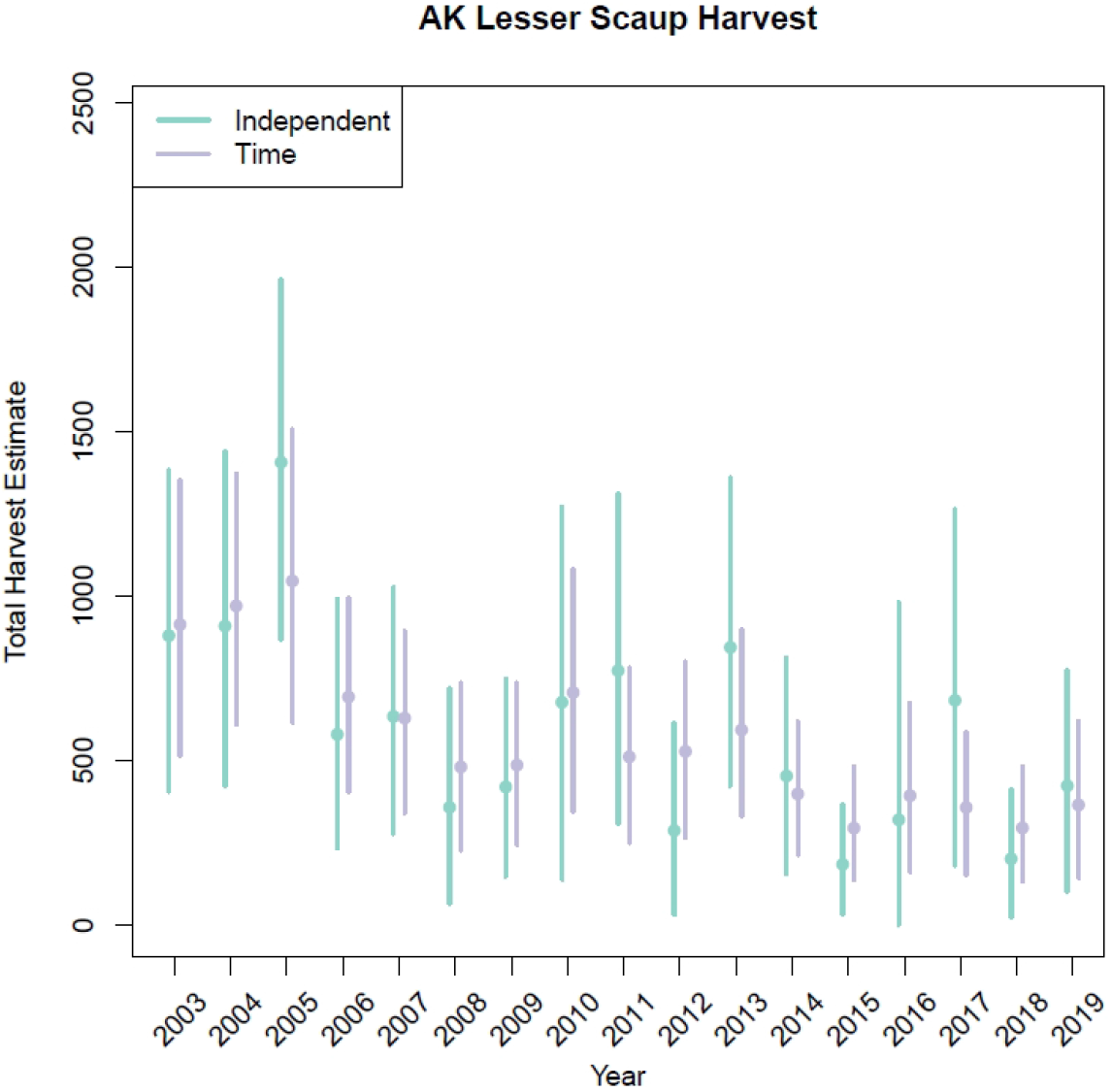
Lesser scaup (*Aythya affinis*) harvest in Alaska (AK) as an example of a species with more temporal correlation, resulting in more shrinkage of the temporal point estimates towards estimates in neighboring years.

**Figure 5:**
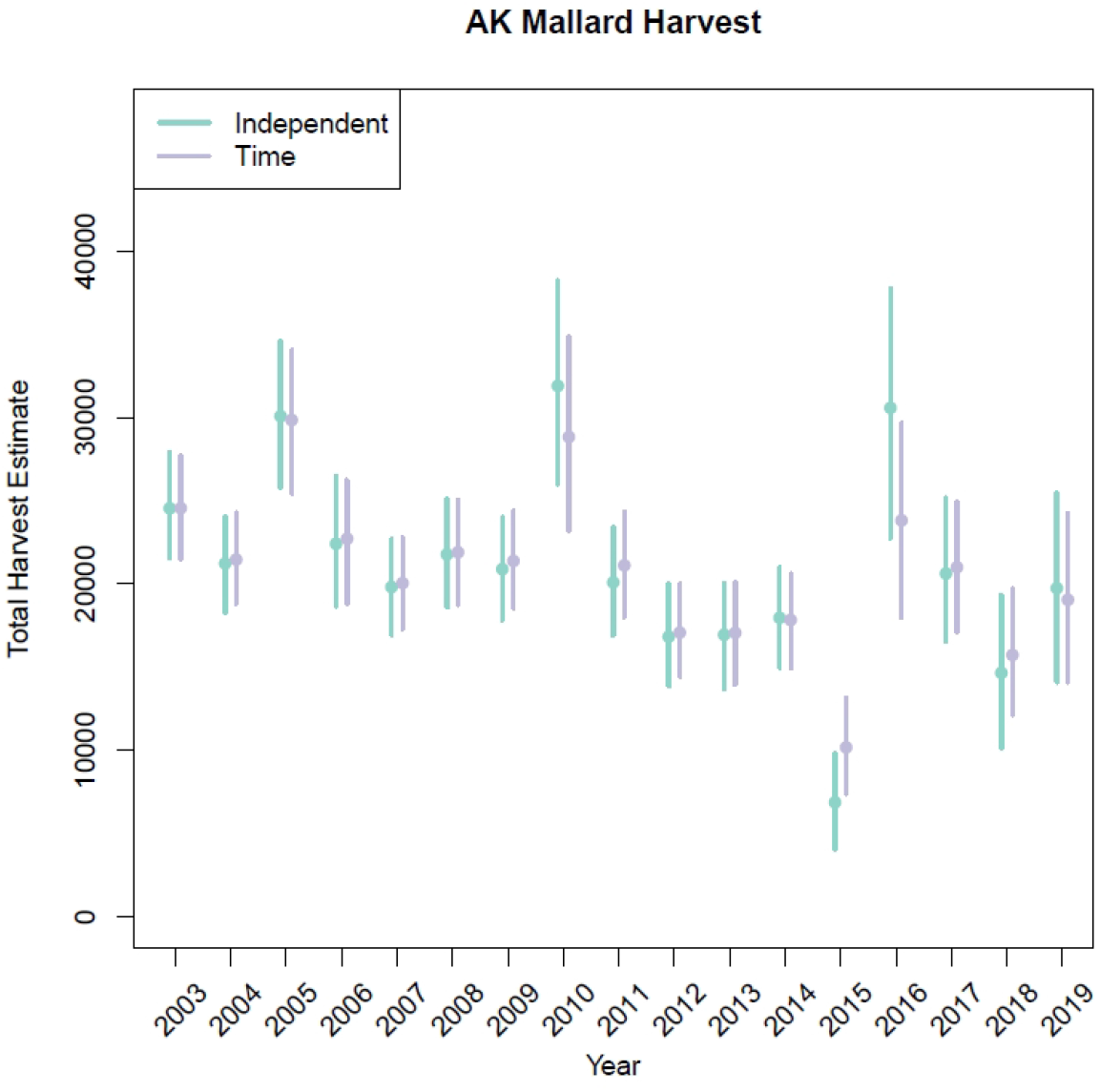
Mallard (*Anas platyrhynchos*) harvest in Alaska (AK) as an example of a species with less temporal correlation, resulting in less shrinkage of the temporal point estimates towards estimates in neighboring years.

**Figure 6:**
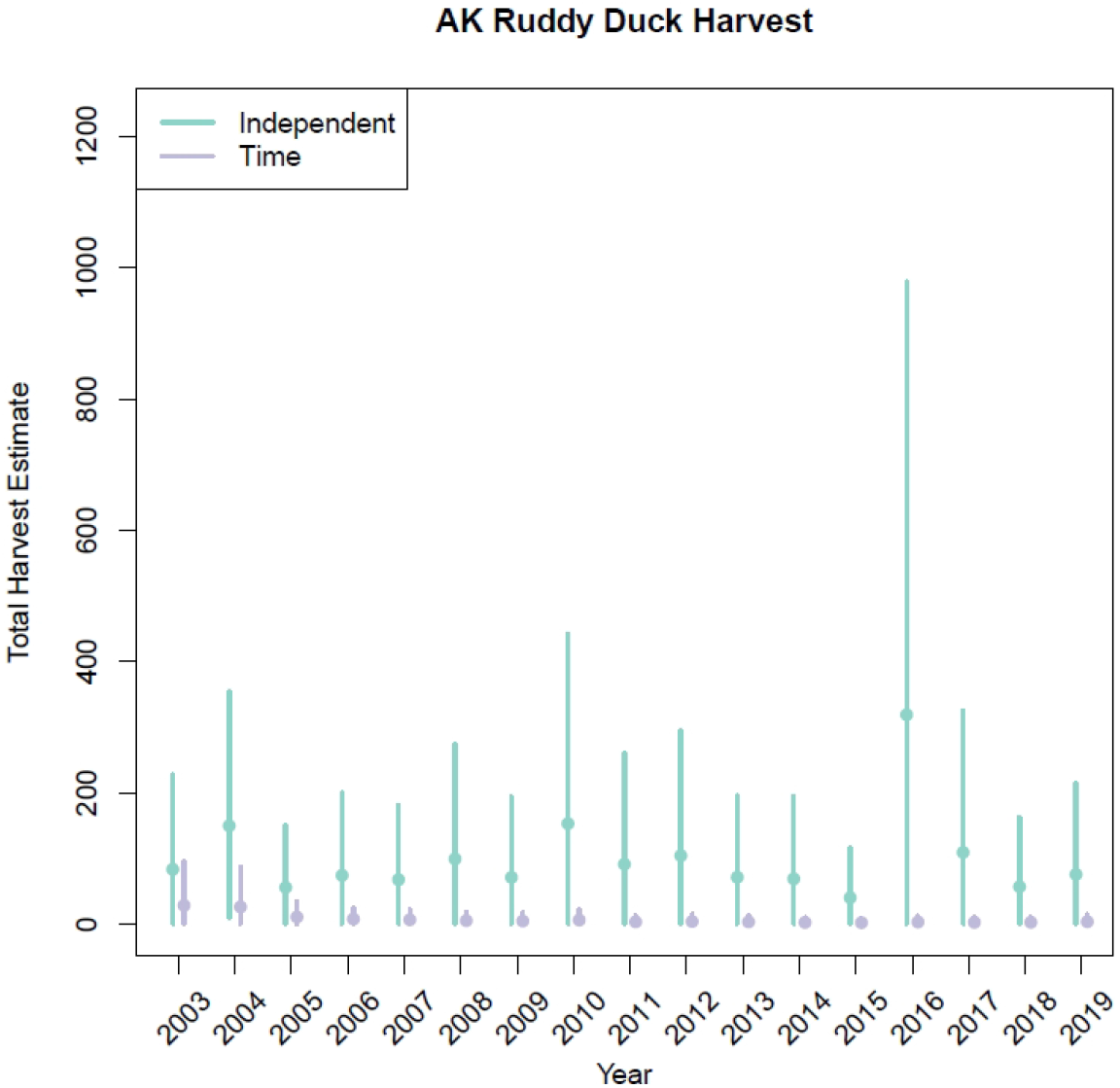
Ruddy duck (*Oxyura jamaicensis*) harvest in Alaska (AK) as an example of a species that is rare in the PCS data. The PCS data from single years are not sufficient to overpower the vague yearly prior distributions with an expected value of equal species proportions, but pooling data over years with a temporal random walk process more appropriately estimates a lower species proportion in the PCS model.

### Results - Flyway-Level

Results for flyway-level estimates (Figure 7) are consistent with those just presented for the state-level analysis (Figures 1-3). The flyway-level point estimates between the independence and time models were largely similar, except for harvest of Brant and some geese, which were estimated somewhat lower in the time model, and posterior standard deviations were smaller in the time model. However, there was more variation in the differences between the CVs at the state level. Figure 8 shows an example of a commonly harvested species where the independence and time model point estimates are similar but are modestly more precise in the time model. Conversely, Figure 9 shows an example of a rarely harvested species where the time model estimates are lower than those from the independence model, and the 95% HPD intervals largely do not overlap those from the independence model.

**Figure 7:**
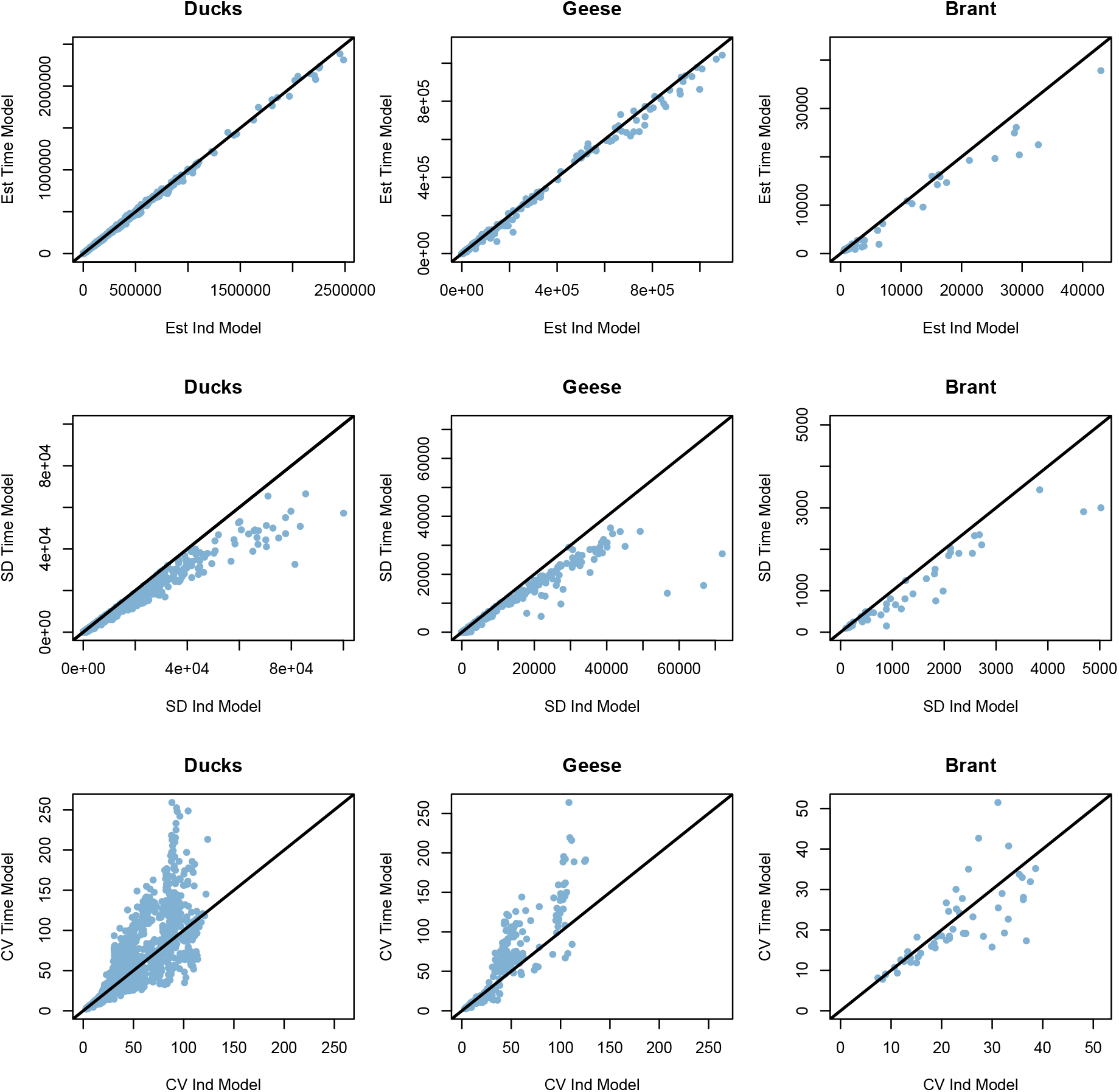
Comparison of flyway by species by year harvest point estimates, posterior standard deviations, and CVs for each species class pooled across species and years. Estimates from the independence and time models are on the x and y axes, respectively. The black line indicates equality between the two models. Note, seaducks and ducks are combined into a single duck category at the flyway level.

**Figure 8:**
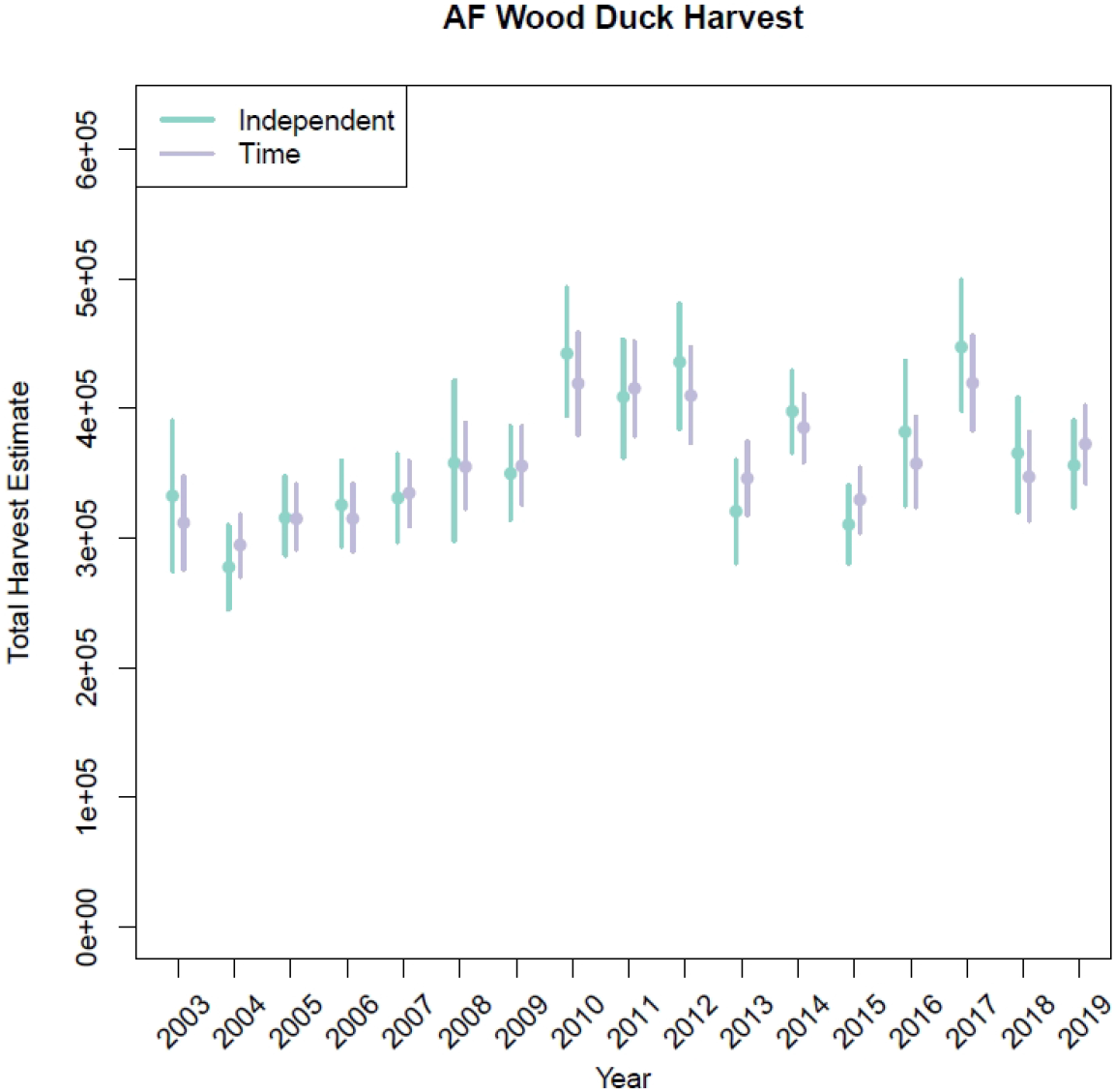
Wood duck (*Aix sponsa*) harvest estimates in the Atlantic flyway (AF) as an example of a species that is commonly harvested, where the time model estimates are similar to the independence model.

**Figure 9:**
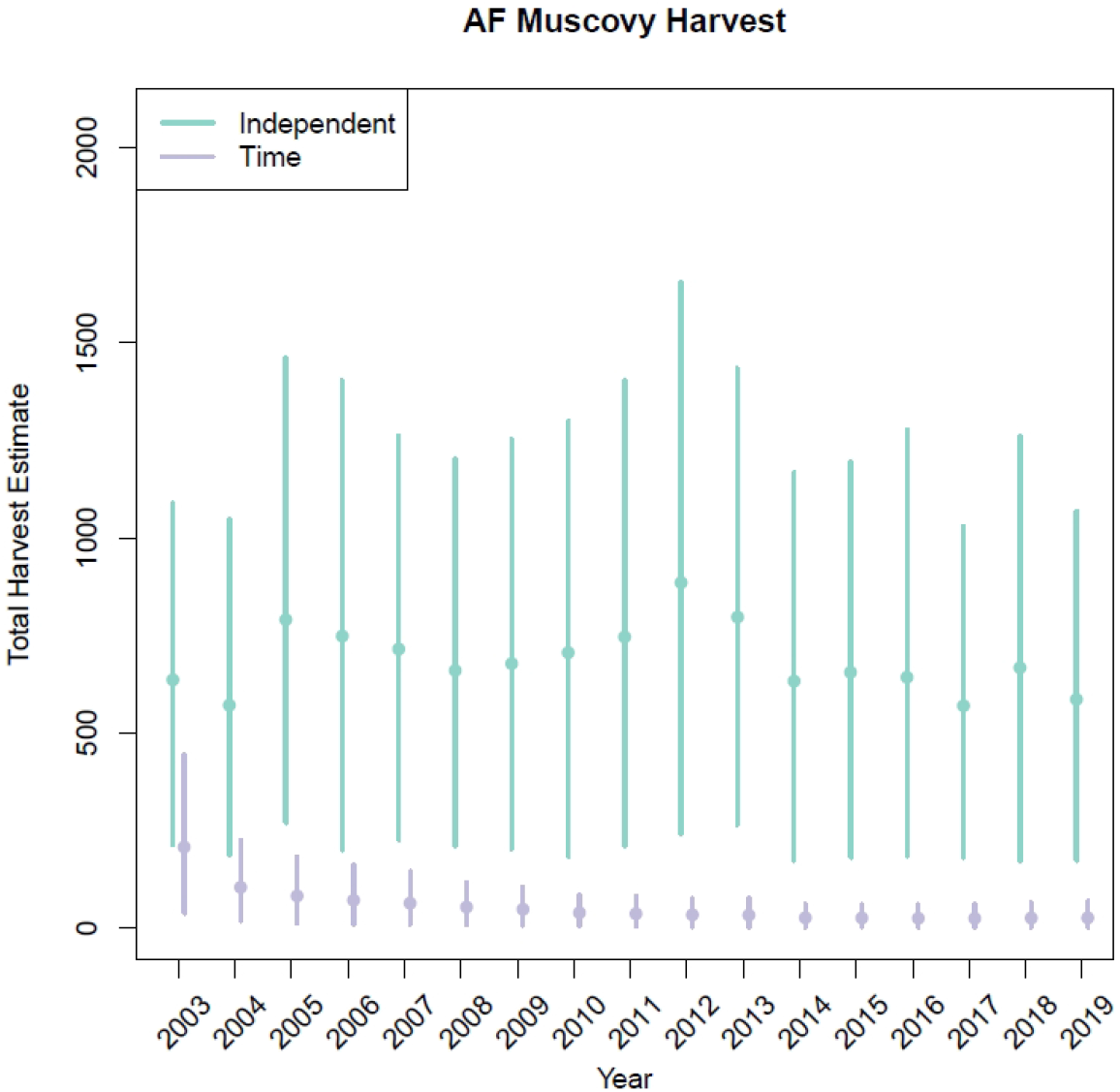
Muscovy (*Cairina moschata*) harvest in the Atlantic flyway (AF) as an example of a species that is rarely harvested, where the time model estimates are much lower than the independence model.

## Discussion

We described and applied two models that integrate data from the Waterfowl Harvest Survey (WHS) and Parts Collection Survey (PCS) data to produce species-specific harvest estimates at the state and flyway levels with the variance of both data sets propagated to the species-specific harvest estimates. The time model that considers temporal correlation in the total harvest and species proportions generally produced more precise parameter estimates for commonly harvested species that are likely to also be more accurate. For more rarely harvested species, the time model does not necessarily increase precision as measured by the CV–both the point and standard deviation estimates are lower in the time model, and the difference in the CV between the time and independence model depends on the relative changes in the point and standard deviation estimates. However, the independence model estimates for rarely harvested species are likely to be systematically overestimated due to the difficulty estimating very small species proportions in the PCS analysis with sparse yearly data sets at the state level–there is not enough information in the data to fully remove the influence of the moderately informative yearly prior distributions on the posterior distributions. By including the temporal correlation in the prior model, the time model is able to pool information across time and reduce the positive bias seen in the independence model estimates. Therefore we conclude that the time model estimates are likely more accurate for both commonly and rarely harvested species, with more appropriate posterior standard deviation and interval estimates.

A limitation of the species and age-specific harvest estimates from these models is the relatively small number of submitted parts of many species and sex classes, leading to highly imprecise estimates of those class proportions. Obtaining larger sample sizes would incur more cost and staff time for processing, making this a challenging problem to deal with. Naturally, this problem might be alleviated in the future if the PCS program adopts an App-based submission framework which would make it simpler for hunters to submit data on harvested birds. The model also assumes that the individual parts from the PCS survey are independent samples of the population of harvested birds. This assumption is likely violated, as individual hunters submit multiple parts. If individual hunters are targeting species, then disproportionate representation by a small number of hunters could induce bias in the parts sample with respect to species and sex, and this bias may vary by state, depending on the focus of individual hunters selected to contribute to the PCS. A potential resolution is modeling hunter-level data, allowing for the species-sex proportions to vary by hunter, and then using a more complex multi-level model to aggregate hunter-level proportions to states and flyways.

## Acknowledgements

We thank for the following USFWS biologists for their support of the project: Kathy Fleming, Joshua Dooley, Tony Roberts and Bob Raftovich. Funding for this project was provided by U.S. Fish and Wildlife Service, Branch of Monitoring and Data Management, the Arctic Goose Joint Venture and the Sea Duck Joint Venture. Any use of trade, firm, or product names is for descriptive purposes only and does not imply endorsement by the U.S. Government.

